# REALDIST: Real-valued protein distance prediction

**DOI:** 10.1101/2020.11.28.402214

**Authors:** Badri Adhikari

## Abstract

Protein structure prediction continues to stand as an unsolved problem in bioinformatics and biomedicine. Deep learning algorithms and the availability of metagenomic sequences have led to the development of new approaches to predict inter-residue distances—the key intermediate step. Different from the recently successful methods which frame the problem as a multi-class classification problem, this article introduces a real-valued distance prediction method REALDIST. Using a representative set of 43 thousand protein chains, a variant of deep ResNet is trained to predict real-valued distance maps. The contacts derived from the real-valued distance maps predicted by this method, on the most difficult CASP13 free-modeling protein datasets, demonstrate a long-range top-L precision of 52%, which is 17% higher than the top CASP13 predictor Raptor-X and slightly higher than the more recent trRosetta method. Similar improvements are observed on the CAMEO ‘hard’ and ‘very hard’ datasets. Three-dimensional (3D) structure prediction guided by real-valued distances reveals that for short proteins the mean accuracy of the 3D models is slightly higher than the top human predictor AlphaFold and server predictor Quark in the CASP13 competition.

## 1 Introduction

With a background of inter-residue contact prediction as the accepted paradigm for decades [1], the field of protein structure prediction is now shifting to the paradigm of predicting the probability of distance intervals, also known as ‘distograms’ [2]. After the introduction of deep learning-based methods [3, 4], predicted contact information arose as a key intermediate step towards accurate ab initio structure prediction. However, after DeepMind and the Xu group demonstrated that predicting the probabilities of distance bins can be more informative than their binary counterparts (contacts) [5], many groups are now pursuing similar approaches. Notable methods such as trRosetta [6], the ResNet/Densenet-based method [7], and DeepDist [8] have shown results similar to or better than the top groups in the CASP13 competition. Overall, the paradigm of distogram prediction is currently “the” promising direction to solve the long-standing problem of protein folding. While multiple sequence alignments and deep learning hold a lot of promise, undeniably, we are far from a convergence on the algorithms and methods for *ab initio* protein structure prediction. Therefore, an exploration of alternative endeavors is needed. As the current methods for contact and distogram prediction mature and become even more accurate, we will naturally ask if we can predict the actual real-valued distances and thereby convey these distances as they are in nature.

This work focuses on a new (third) paradigm—one of predicting real-valued distances, i.e. predicting what the distances truly are. Contacts or distograms, on the other hand, are human defined zero-one or multi-class labels [9]. After Kukic et al. [10] and Walsh et al. [11] introduced the idea of real-valued distance prediction, the author of this work reintroduced this paradigm in the context of deep learning and recently released an open-source framework for distance prediction [12]. In this work, we demonstrate that real-valued distance prediction alone can deliver on par or better precision compared to the state-of-the-art contact and distance prediction methods. Towards demonstrating that real-valued distance prediction is a direction full of promises, this work establishes a groundwork.

## 2 METHODS

### 2.1 Development set

Current approaches to training a deep learning model either use a smaller set of a sequence-similarity-reduced database such as PISCES [13] or a structural-similarity-reduced database such as CATH [14]. For example, methods such as Raptor-X [4] use a version of a sequence-similarity-reduced dataset, whereas more recent methods such as AlphaFold [2] use a version of a structural-similarity-reduced dataset. A recent work [15] suggests that future methods should focus on the use of structural-similarity-reduced datasets such as CATH [14] or ECOD [16] for training and evaluating deep learning methods. While sequence-similarity-reduced databases ensure the representation of the known protein sequences, the structural similarity reduced datasets ensure the representation of the structural fold space.

In this work, we were interested in training a deep learning model that learns from both the protein sequence space and the structural space. To achieve this, we merged a version of the PISCES set and a CATH set. We used the May 2018 release of the PISCES dataset having 27,832 chains, curated using the following parameters: percentage identity cutoff = 70%, resolution cutoff = 3.0Å, R-factor cutoff = 1.0, and X-RAY structures excluded. The dataset is maintained by the Dunbrack Lab and is available at http://dunbrack.fccc.edu/. We further cleaned this list by removing chains that had large structural gaps after the removal of non-standard amino acids or if they had lesser than 12 residues. Chains longer than 512 residues were trimmed by keeping only the first 512 residues. Our final PISCES set included 27,319 chains (set *P*). Similarly, we cleaned the CATH v4.2 domains (released in April 2018) consisting of 31,289 structural domains, to obtain a final set consisting of 24,864 unique chains (set *C*). The CATH dataset is available at https://www.cathdb.info/. Finally, we merged the two sets to obtain a total of 43,071 unique protein chains (*P*∪*C*). This final development set is less than the sum of the two because of a large number of overlapping protein chains. A random set of 200 chains from the development set were selected as a validation set, leaving the remaining chains as the training set. Notably, our development set consists of proteins released before May 2018, allowing us to evaluate our method on targets released afterwards, such as the CASP13 targets.

### 2.2 Deep learning features

Successful contact prediction methods such as ResTripLet and TripletRes [17] demonstrate that precision matrix and covariance matrix are the key features that drive contact precision. In this work, we use six input feature sets: (1) a reduced precision matrix (231 2D features), (2) a position specific scoring matrix along with the sum (22 1D features) obtained using a script in the trRosetta method, (3) composition of the amino acids (20 1D features), (4) entropy (one 1D feature) obtained using the script from trRosetta, (5) CCMpred [18] and FreeContact [19] predictions (two 2D channels), and (6) potential (one 2D channel). After translating each one-dimensional (1D) feature into two two-dimensional (2D) features—by tiling and transposed tiling - we have a total of 320 2D channels. An inverse covariance matrix, also known as the precision matrix, typically consists of 21 × 21 (= 441) channels, where 21 refers to the 20 standard amino acids and a gap character. For a protein of length L, a precision matrix P is a L x L x 21 × 21 matrix, where each 21 × 21 matrix captures the direct coupling correlations for 21 × 21 residue type pairs at the corresponding residue pair position. We tweaked the process of generating a precision matrix discussed in ResPRE [20] to build a half-compressed version, with only 231 channels, so our feature files for the entire development set can be fit in a 2 Terabyte solid state disk (SSD). Wu et al. have discussed this technique of compressing the relationship among variables for a standard covariance matrix [21]. Here we extend the idea to compress a precision matrix instead of a covariance matrix. Specifically, if *i* and *j* correspond to the pairing residues, we obtain the reduced precision matrix *P*_*reduced*_ using Eq(1).

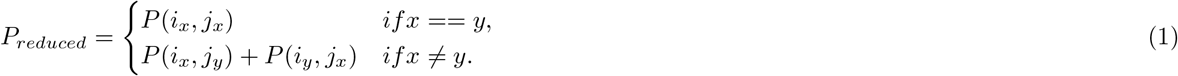

### 2.3 Deep learning setup

Our network architecture is a variant of a standard residual network (ResNet). As shown in **Figure 1**, each residual block in our network consists of a batch normalization layer followed by an exponential linear unit (ELU) activation, a 2D convolution layer consisting of 128 3 × 3 filters, a dropout layer with a dropout rate of 20% followed by ELU activation, and finally a 2D convolution layer consisting of 128 filters that alternate between 3 x 3 and 1 × 5 kernels as well as alternating dilations of 1, 2, and 4. In addition to the 128 residual blocks, the architecture has a 2D convolutional block to shrink the input volume (128 × 128 × 322) so the ResNet block receives a 128-channel input and a 2D convolutional block that receives the output of the ResNet block and shrinks the number of channels to one, effectively predicting real-valued distances. With 128 residual blocks, and effectively 256+ convolutional layers, the model has 29.5 million parameters. We train our model at a fixed window of 128 x 128. In other words, in each model training/validation task, we only predict the distances between two sequence pairs, each of which is a maximum of 128 residues long. It is intriguing that such a setting allows the model to learn the distances anywhere on the distance map for a protein of any length. With a batch size set to two, one epoch of training takes about 8 hours in a TITAN RTX GPU when the features and distance maps are all loaded from solid state disks.

**Figure 1:**
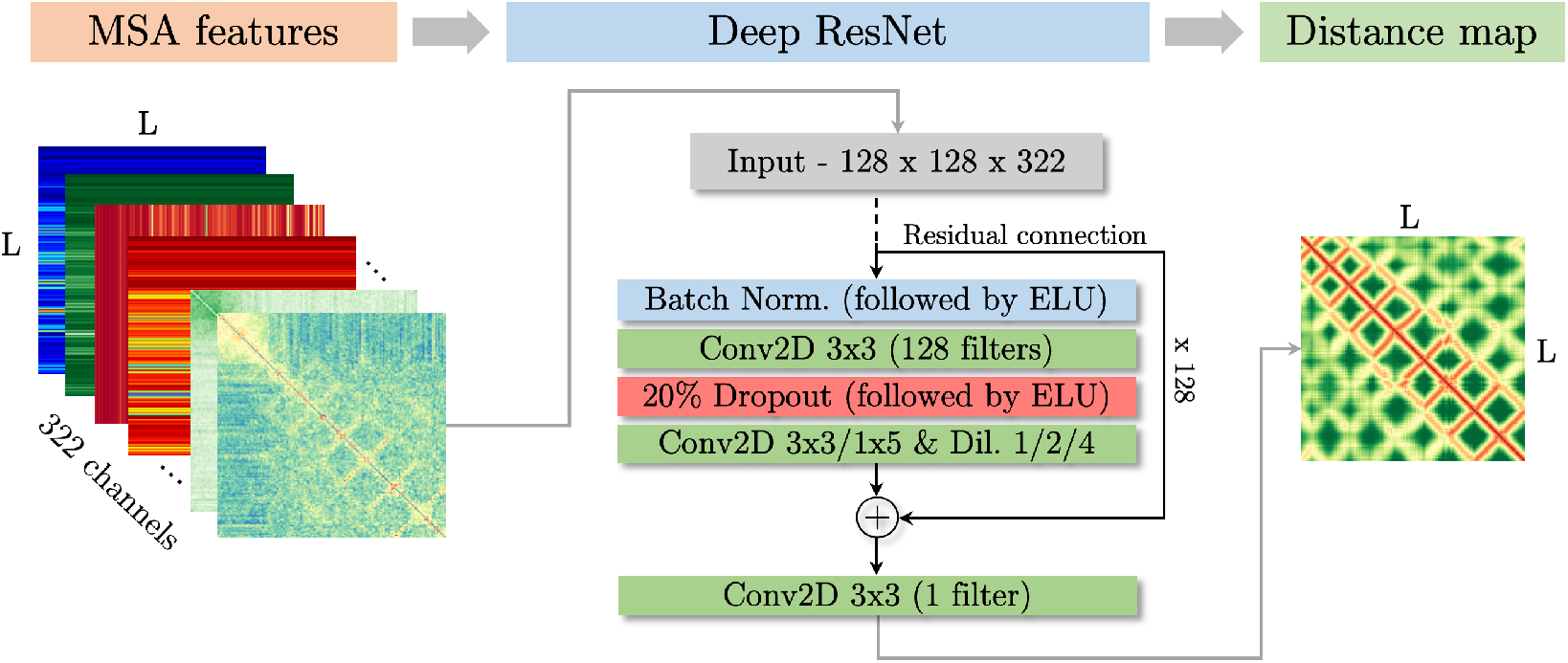
Three major steps involved in real-valued distance prediction: obtain input volume from multiple sequence alignment (MSA) features, deep ResNet training, and real-valued distance map prediction.

To effectively predict real-valued distances, the biggest challenge is the design of a right loss function. Commonly used loss functions automatically focus on long physical distances in the distance map (because loss is higher for longer distances), which are difficult to predict and also less informative about the interactions in the structure. In a recent work, we demonstrated that one approach to attack this problem is to reciprocate the distances before feeding them to the deep learning model [12]. It is important to note that our approach differs completely when compared to the DeepDist approach, where Wu et al. [8] have discussed using the mean squared error (MSE) loss and flooring distances to 16Å, i.e. *d*[*d >* 16] = 16. Subsequently, we investigated alternative methods to reciprocate the distances so the contacts derived from the predicted distances are at least as good as when only contacts are predicted [22]. We were successful in doing so using a special distance transformation function. Specifically, if *d*_*ij*_ is a true distance between residues *i* and *j*, we transform the distance into a new reciprocated distance 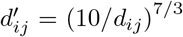. A deep learning model’s output distances are then recovered using 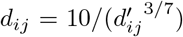. Notably, this is label engineering and not feature engineering. Reciprocating distances and recovering, enables the use of any standard regression loss function. We found the best performance to come from the logarithmic cos hyperbolic loss function and the ‘rmsprop’ optimization algorithm.

### 2.4 Generating multiple sequence alignments

For the chains in our development set, we generated multiple sequence alignments (MSAs) using the ‘HHBlits’ program in the HHSearch package [23] with the ‘Uniclust30’ sequence database [24] released in October, 2017 (e-value = 0.001, iterations = 3, and coverage = 40%). For difficult (free-modeling) protein sequences, where the standard methods such as HHsearch or PSIBLAST [25] fail to generate a reasonable number of alignments, the use of Metagenomic sequence databases have been proposed [21]. Hence, for the protein sequences in our test sets (i.e., CASP13 and CAMEO targets) we generated MSAs using DeepMSA [26]. We augmented DeepMSA using metagenomic sequence databases from multiple sources (see **Table S1**). These databases are large in size, ranging from 50GB to 450GB when uncompressed. Since, running DeepMSA with these databases is slow with conventional hard-drives, we used solid state disks (SSDs). Even with SSDs, our alignment generation with DeepMSA takes about up to two hours for a sequence of around 250 residues. For accurate distance prediction, although a pre-trained deep learning model can be run on a regular GPU (or even a CPU), high quality alignment generation for difficult protein sequences comes with high computational costs (high random-access memory (RAM) and large SSDs).

The common parameters such as input sequence coverage and ‘e-value’ for running alignment prediction tools (JackHmmer and Hhblits) are ineffective when the input sequence is long. Typical coverage parameter values such as 60% are not effective when we are searching for alignments for a long sequence. This is because the sequence hits for some sub-sequences of our input sequence with lengths that were too short to meet the coverage parameter are not reported. Although structural domain prediction is a possible route to explore, previous CASP participants have reported that a failed domain splitting can severely affect precision [27, 28]. In this work, we evenly split input sequences longer than 256 residues into overlapping sub-sequences of 256 residues, with an overlap of 128 residues. For example, a 500-residue-long protein will have three subsequences: (1) 1 to 256 residues, (2) 128 to 384 residues, and (3) 244 to 500 residues. Then, we generate MSAs for all the sub-sequences including the original full input sequence. We found that merging these MSAs is not a straightforward problem. Hence, we predict distances using the MSAs for each sub-sequence and merge the overlapping distance maps by selecting the minimum predicted distance at each pixel. Selecting smallest distance (short physical distance and not necessarily short sequence separation) works well in the context of the REALDIST method because REALDIST is designed to predict short distances more accurately, i.e., loss is higher for short distances.

### 2.5 Evaluation of predicted distances

In the absence of an established metric to evaluate predicted real-valued distances, we chose to first evaluate (a) the precision of contacts derived from the predicted distances using precision metrics [29, 30], which has been the choice of other successful groups and the CASP organizers, and (b) distances using various metrics obtained using DISTEVAL [31], a recently developed distance evaluation tool available at http://deep.cs.umsl.edu/disteval/. To obtain contacts from predicted distance maps, we assign contact scores wherein the shorter predicted distances have higher scores than the longer ones and a score of 0.5 is assigned for a predicted distance of 8Å. Therefore, if *d*_*ij*_ is the predicted distance between two residues, *i* and *j*, then the corresponding confidence score *p*_*ij*_ is,

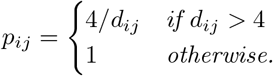

Since real-valued distance prediction is a recently uncovered promise, it remains to be explored what the most effective ways of evaluating predicted distances are. While the standard metrics for calculating distance errors such as mean absolute error (MAE), root mean squared error (RMSE), and local distance difference test (lDDT) [32] scores may be obvious choices, multiple metrics may complement each other’s strengths and weaknesses. We sought to evaluate the real-valued distances predicted by REALDIST using MAE, *Cβ*-lDDT score, and Pearson correlation coefficient (PCC) and compare with other distance prediction methods. To calculate MAE and Pearson correlation coefficient, all distances predicted within 15Å and with minimum sequence separation of 12 residues are considered, i.e., medium and long-range distances are evaluated. Similarly, for calculating *Cβ*-lDDT, only carbon-beta atoms are considered (*Cα* in case of glycine), and the ‘R’ parameter is set to the default value of 15Å. Also, for comparsion with the predictions by the trRosetta method, the ‘distograms’ are translated to real-valued distance maps by ‘flattening’, i.e, we chose the center of the distance bin with the highest probability as the real-valued prediction. For an even more rigorous evaluation of predicted distances, we built 3D models using all the predicted real-valued distances below 12Å using the CONFOLD tool and selected the top-one model, out of 20, for evaluation. Since CONFOLD does not rely on any template or template derived information, it is an ultimate assessment of predicted distance maps. The top models (not the best models) are evaluated using TM-score, RMSD, and GDT-TS [33].

### 2.6 Structure prediction from real-valued distance maps

We translated the predicted real-valued distances to upper and lower bounds to generate restraints for 3D modeling using the Rosetta *ab initio* protocol. For a predicted real-valued distance *d*, we calculated error range, *δ* = 0.03\**d*\**d*, where *d* is the predicted real-valued distance; the lower bound *l* = *d* — *δ*/2, and the upper bound *u* = *d* + *δ/*2. This empirical rule of setting a higher error range for longer distances follows the design of our loss function, which focuses on learning to predict shorter (not necessarily short-range) distance values before longer ones; therefore, shorter distance predictions are more likely to be correct. With these restraints, we built 1000 decoys using the Rosetta ab initio protocol and selected the model with the lowest energy score as the predicted model. We assert that the methods for calculating *δ* can be further optimized for a chosen 3D modeling protocol or even be predicted as an additional output channel of a deep learning model. For building models using CONFOLD, however, we used all predicted distances up to 12Å without constraint relaxations.

## 3 RESULTS

In addition to performing some of our own evaluation and ablation studies, here the author of this work (referred as ‘we’ hereafter) rigorously evaluates REALDIST by comparing its performance with some of the most successful state-of-the-art methods that are publicly available, using the most difficult datasets. Specifically, we focus our evaluation on three datasets: a) 131 hard targets from the Continuous Automated Model Evaluation (CAMEO) challenge [34] released between December 8, 2018 and June 1, 2019, b) a subset of 66 ‘very’ hard targets from the set of 131 hard targets, and c) 25 CASP13 free-modeling (FM) targets released in 2018 (31 domains in total). These datasets are also used by other state-of-the-art methods such as trRosetta [6] and DeepDist [8] to evaluate the precision of their methods. Since the REALDIST pipeline (see **Methods**) does not rely on any databases curated before May 2018 for its training, it can be safely assumed that none of the targets in these sets were used in deep-learning model training. The metagenomic sequence databases used in DeepMSA [26] to generate multiple sequence alignments were also curated before May 2018.

### 3.1 Evaluation of contacts derived from predicted real-valued distances

On the 25 free-modeling (FM) targets released in the CASP13 competition, we tested the accuracy of REALDIST. Following the well-known procedure established by the CASP organizers, during the prediction step, we predicted real-valued distance maps for the entire sequence of the CASP targets without any knowledge of the structural domain boundaries. Next, contact probabilities were obtained from the real-valued distance prediction matrix by using a 4*/d* rule (see **Methods**) and these contact maps were evaluated using the ‘precision’ metric [29, 30]. Evaluation of the top L/5, and top L long-range contacts, as summarized in **Table 1**, reveals that REALDIST considerably outperformed the top CASP13 performers. Here, L stands for the number of residues in the true structure. While the top CASP13 performer (Raptor-X, Group #498) scored a precision of 70.2% (top L/5) and 44.7% (top L), the precision values of the contacts obtained from REALDIST predictions were 79.0% (top L/5) and 52.2% (top L). REALDIST’s better precision on the top L metric, which is a much bigger set than the top L/5, also demonstrated that REALDIST does not just predict a few portions of the true contacts correctly. An example of our method’s improved performance is shown in **Figure 2** for target T0968s1 wherein the precision of the top L long-range contacts is 55.1%, as predicted by REALDIST, while the precision values of the top three methods are: Group #498 (35.6%), Group #032 (31.4%), and Group #180 (28.8%).

**Table 1:** Precision of top L/5 and top L contacts predicted by REALDIST, the top three contact prediction groups for the 31 CASP13 free-modeling domains, and trRosetta on CASP13 free modeling targets and CAMEO hard targets. Similar to REALDIST, trRosetta’s baseline method is a model trained without MSA subsampling, MSA selection, or orientation prediction. Long-range contacts (*s* ≥ 24) and medium- and long-range contacts (*s* ≥ 12) are evaluated separately.

| Dataset | Method | $s \geq 24$ | | $s \geq 12$ | |
| --- | --- | --- | --- | --- | --- |
| | | $P_{L/5}$ | $P_L$ | $P_{L/5}$ | $P_L$ |
| 31 CASP13<br>FM domains | CASP13 Group #498 | 70.2 | 44.7 | 87.3 | 61.3 |
|  | CASP13 Group #032 | 65.5 | 42.4 | 84.7 | 61.0 |
|  | CASP13 Group #180 | 64.3 | 41.5 | 84.8 | 58.8 |
|  | trRosetta baseline | 68.6 | 44.3 | 80.0 | 60.7 |
|  | trRosetta | 78.5 | 51.9 | **89.9** | **70.2** |
|  | REALDIST with trRosetta’s alignments | 72.9 | 50.1 | 86.8 | 67.6 |
|  | REALDIST with DeepMSA alignments | **79.0** | **52.2** | 86.5 | 69.7 |
| 131 CAMEO<br>hard targets | trRosetta baseline<sup>a</sup> | 78.1 | 52.1 | 80.1 | 60.4 |
|  | trRosetta | 82.7 | 57.0 | 82.7 | 64.4 |
|  | REALDIST with trRosetta’s alignments | 81.2 | 57.9 | 83.9 | 67.4 |
|  | REALDIST with DeepMSA alignments | **83.8** | **58.2** | **86.4** | **67.7** |
| 66 CAMEO<br>very hard<br>targets | trRosetta baseline<sup>a</sup> | 67.7 | 41.6 | 76.9 | 57.5 |
|  | trRosetta | 75.4 | 48.0 | **81.5** | **62.8** |
|  | REALDIST with trRosetta’s alignments | 72.0 | 48.0 | 75.1 | 57.6 |
|  | REALDIST with DeepMSA alignments | **76.9** | **49.5** | 80.1 | 59.6 |

<sup>a</sup>Trained without MSA subsampling, MSA selection, or orientation prediction.
L denotes the length of the protein sequence.
s denotes minimum sequence separation.

**Figure 2:**
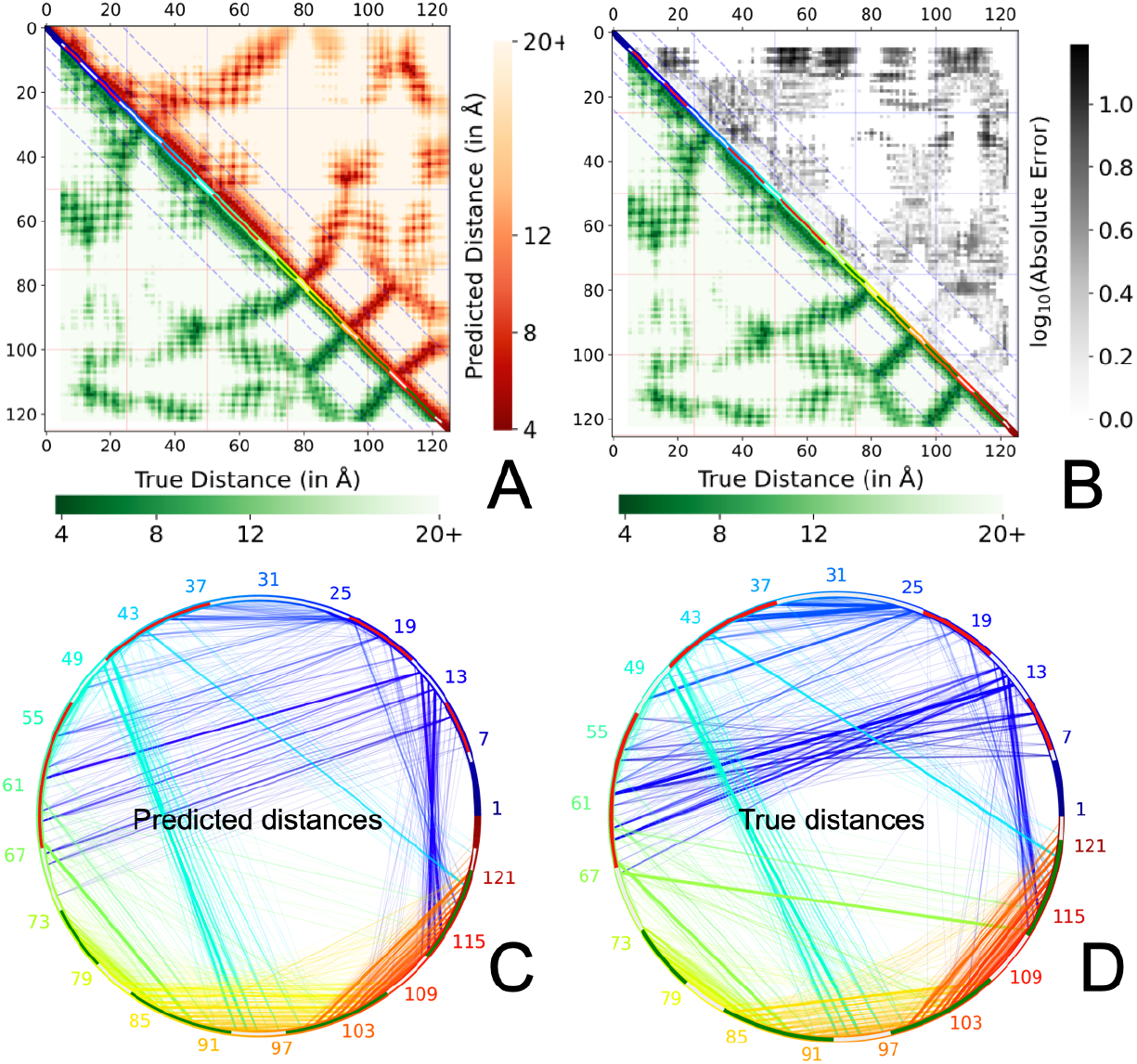
REALDIST’s distance prediction for the CASP13 FM target T0968s1. When the top L/5 and top L long-range contacts obtained from corresponding predicted distances are evaluated, the precision values are 79.2% and 55.1% respectively. (A) The predicted distance map in the upper triangle is compared with the native distance map in the lower triangle. Diagonal lines are the markers of short-range, medium-range, and long-range distances. (B) Absolute error in the predicted distance map highlight the regions missed by the predicted distance map. The circular chord diagrams represent the (C) predicted distances and (D) the native distances, wherein most long-range interactions are captured by REALDIST. Red and green arch regions in the circumference of the chord diagrams correspond to the helix and strand residues in the native structure.

To conclude our distance-derived contact evaluation, we compared the accuracy of REALDIST with trRosetta, a method demonstrated to outperform all CASP13 methods on the 31 free-modeling (FM) target domains from CASP13. trRosetta is also demonstrated to outperform all other methods on the 131 hard targets released in CAMEO. We derived contact probabilities from the trRosetta’s distogram predictions by summing the bin probabilities up to the 8Å bin. Our results, summarized in **Table 1**, demonstrate that distance predictions by REALDIST are appreciably more precise than the trRosetta’s baseline model—top L long-range precision of 52.2% vs. 44.3% on the CASP13 set. REALDIST is comparable to the trRosetta’s baseline method (and not the full version) because similar to REALDIST the baseline model is trained without multiple sequence alignment (MSA) subsampling, MSA selection, or orientation prediction. We also compared REALDIST’s performance with the final trRosetta method (the full version). Results summarized in **Table 1** show that REALDIST performs similar to trRosetta on the CASP13 FM dataset; the precision of the top L long-range contacts was 52.2% for REALDIST and 51.6% for trRosetta. On the 131 CAMEO hard dataset, however, the REALDIST predictions were slightly better; here, the precision of top L long-range contacts was 58.2% for REALDIST and 56.3% for trRosetta (**Table 1** and **Figure 3A**). We further compared the performance using the subset of the CAMEO ‘very’ hard set consisting of 66 harder chains and found that REALDIST outperformed trRosetta when long-range contacts are evaluated.

**Figure 3:**
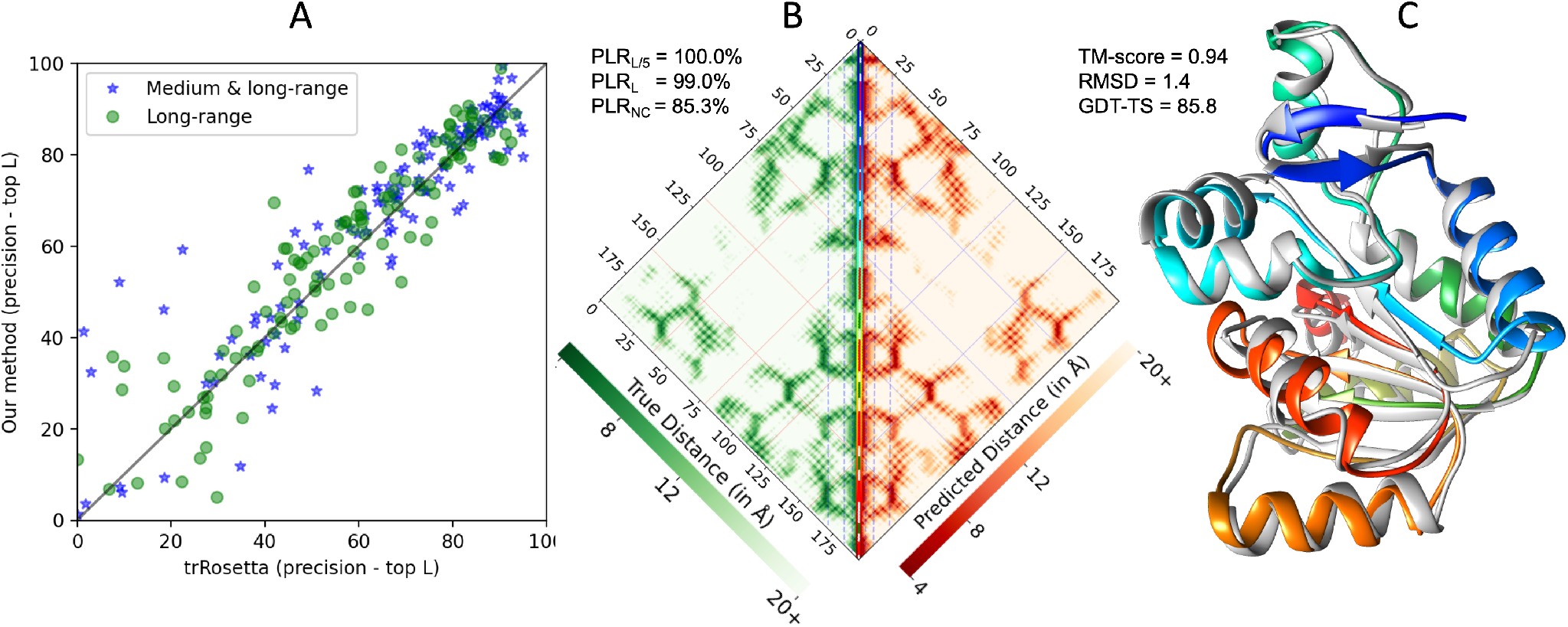
Comparison and evaluation of predicted distances for the CAMEO set. (A) Comparison of the precision of top-L contacts derived from distances predicted by REALDIST and trRosetta on the 131 CAMEO hard set of protein chains, (B) native distance map (in green on the left) and the predicted distance map (on the right) for the CAMEO target ‘5WB4 H’, and (C) native structure (in gray) and the top predicted model (with a multi-color theme) for the CAMEO target ‘5WB4 H’.

### 3.2 Evaluation of predicted real-valued distances

**Table 2** summarizes the evaluation of distance predictions by REALDIST, trRosetta [6], DeepDist [8], and GAN-based method [9] on the 31 CASP13 free-modeling domains. On average, evaluation using the 3D model evaluation metrics (TM-score, RMSD, and GDT-TS) show that REALDIST outperforms all three methods. The generative adversarial network (GAN)-based real-valued distance prediction method [9], to the best of our knowledge, is the only deep learning effort designed to solely predict real-valued distances. The GAN-based method mainly focused on evaluating the predicted real-valued distances by assessing their utility towards building 3D models. On the entire dataset of CASP13 targets (which includes template-based as well as template-free targets), authors demonstrate that real-valued distance prediction using GAN can be remarkably more informative towards accurate structure prediction. As acknowledged by the authors, a key limitation of their work is that the predictions are not blind, i.e., they use structural domain information and make predictions only for the domains. Additionally, since the evaluation does not focus on the free-modeling targets it is unclear how effective their method is for such difficult proteins. To obtain predicted distances from the GAN-based method, we ran the method locally using the alignments generated by the trRosetta method. As summarized in **Table 2**, when distances are predicted using the same input alignments, REALDIST remarkably outperforms the GAN-based method, on the most difficult CASP13 free-modeling dataset.

**Table 2:** Evaluation of predicted real-valued distance maps by DeepDist, trRosetta, GAN method, and REALDIST using contact metrics, distance evaluation metrics (local distance difference test, mean absolute error, and Pearson correlation coefficient), and the accuracy of top one model reconstructed using CONFOLD.

| Method | Contacts |  | Real-valued distances |  |  | 3D models |  |  |
| --- | --- | --- | --- | --- | --- | --- | --- | --- |
| | P<sub>L/5</sub> | P<sub>L</sub> | *C*β-IDDT | MAE | PCC | TM-score | RMSD | GDT-TS |
| DeepDist<sup>a</sup> | 74.8 | 45.9 | **0.52** | 4.1 | 0.54 | 0.36 | 12.4 | 29.1 |
| trRosetta | **79.5** | 51.9 | 0.35 | **2.0** | **0.65** | 0.43 | 11.8 | 36.3 |
| GAN<sup>b</sup> | 57.5 | 37.1 | 0.37 | 4.5 | 0.40 | 0.23 | 15.4 | 16.9 |
| REALDIST | 79.0 | **52.1** | 0.49 | 3.2 | 0.56 | **0.46** | **10.6** | **39.7** |

<sup>a</sup>Distance map predictions downloaded from https://github.com/multicom-toolbox/deepdist
<sup>b</sup>Best of two GAN models obtained from the authors
P<sub>L/5</sub> and P<sub>L</sub> are precision of long-range contacts
*C* $\beta$ -IDDT is calculate with ‘R’ = 15Å and sequence separation = 6
MAE and PCC are calculated with sequence separation $\geq 12$ and for $d_{pred} < 15\text{\AA}$

In **Table 2**, we observe that even though the TM-score and GDT-TS of the top-one models with REALDIST distance predictions are higher than those of other methods, the MAE is quite high and Pearson correlation coefficient is quite low compared to the trRosetta method (MAE 2Å for trRosetta vs 3.2Å for REALDIST). These evaluation results contradict with the accuracy of the reconstructed top-one 3D models. Upon investigating, we found that the metrics such as precision, MAE of predicted distances below a certain threshold, or PCC of predicted distances only assess the predictions with the predictions as the reference, i.e., if the prediction method predicts fewer distances below the chosen threshold (compared to the number of true distances), and the predictions are correct, then metrics favor such methods. In contrast, metrics such as *Cβ*-lDDT score evaluate prediction with the native as the reference. The trRosetta method, in particular, has the *Cβ*-lDDT score considerably lower than REALDIST and DeepDist methods. By comparing the proportion of the number of true distances below 12Å with the predicted distances below 12Å, for the 31 domains in the CASP13 FM dataset, we found that the trRosetta method is a conservative predictor, i.e., prefers to predict rather fewer but accurate predictions (see **Figure 4**). This approach, as it turns out, misses some important distance interaction hubs (i.e., short physical distance patches in the distance map with long-range sequence separation) because of lower coverage, as revealed in the evaluation of the reconstructed 3D models. **Figure 5** illustrates this coverage issue with the CASP13 FM target T0950 as an example.

**Figure 4:**
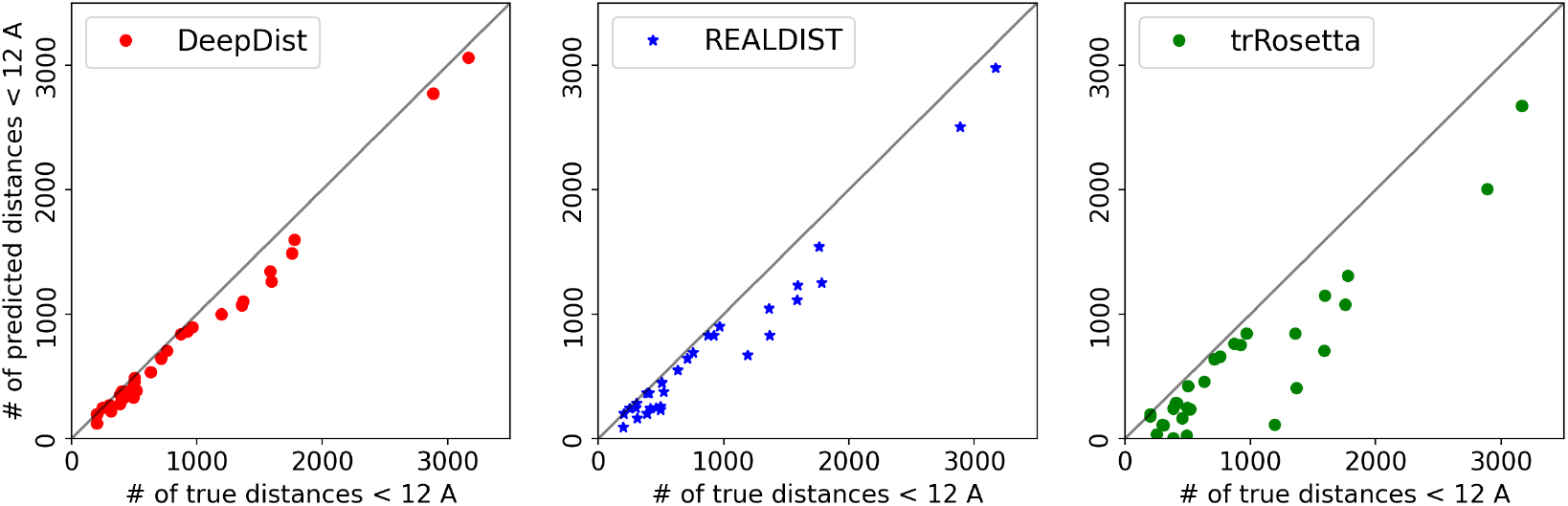
Number of true versus predicted distances below 12Å by DeepDist, REALDIST, and trRosetta for the 31 domains in the CASP13 FM set. Although this for this plot we chose distances below 12Å (the threshold used for reconstruction using CONFOLD), a similar trend is observed for the 15Å threshold.

**Figure 5:**
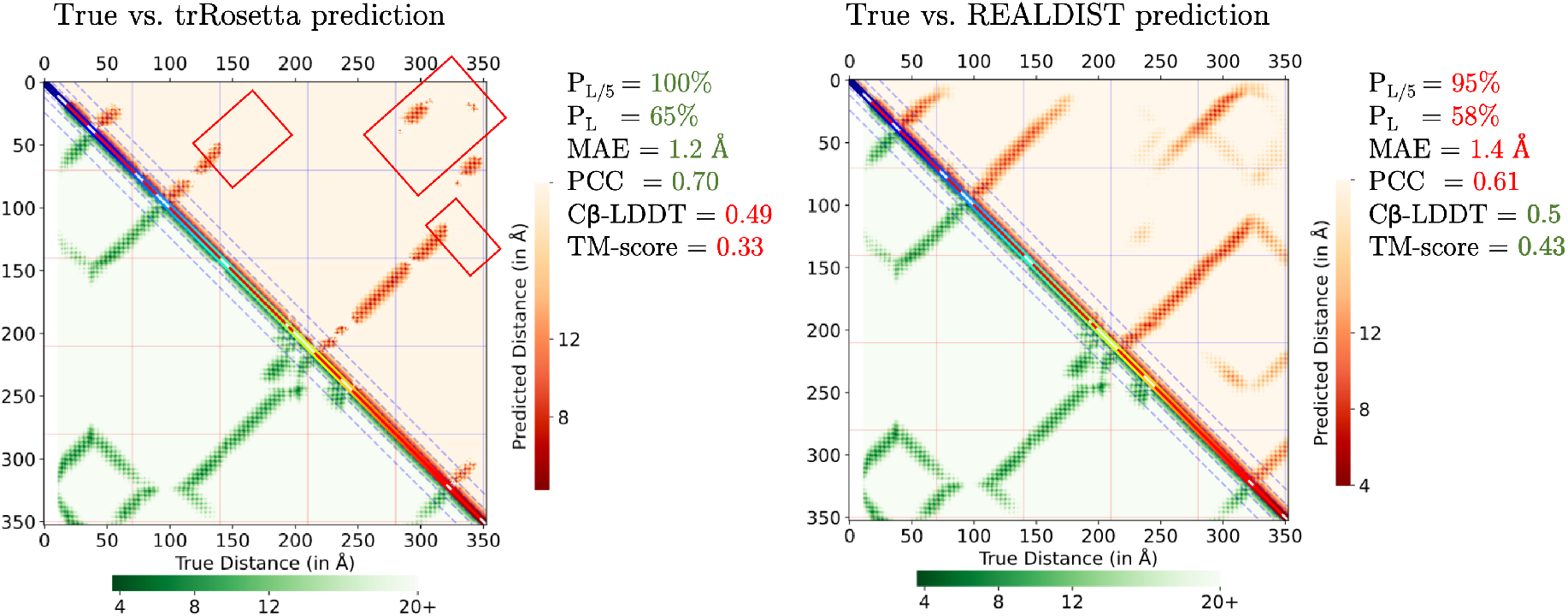
Distance predictions for the CASP13 FM target T0950 by trRosetta and REALDIST evaluated using precision of top L/5 and top L long-range contacts (P_L/5_ and P_L_), mean absolute error (MAE) and Pearson correlation coefficient (PCC) of distances predicted below 15Å, *Cβ*-lDDT of non-local distances (*s*≥6), and TM-score of the top one model reconstructed using CONFOLD. trRosetta misses some ‘short distance’ regions (red square boxes) the prediction appears more accurate when evaluated using P_L/5_, P_L_, MAE, and PCC metrics. However, this low coverage is assessed by *Cβ*-lDDT score and the TM-score of the top one model.

### 3.3 Which contributes more, deep learning or sequence alignments

REALDIST’s accuracy may come from (a) the deep learning model trained on the large number of protein chains, and/or (b) multiple sequence alignments generated using DeepMSA [26]. To investigate if the improvement is not just from the alignments we generated, we switched the input alignments with the alignments by the trRosetta method and predicted distances for the CASP 13 FM dataset and the CAMEO datasets. Only a slight decrease in performance (around 2 percentage points) was observed for both the CASP13 set and the much larger CAMEO dataset (see **Table 1**). This suggests that the improved performance of REALDIST comes from both the deep learning model and the deep and high-quality alignments. Notably, unlike trRosetta, our deep learning method predicts real-valued distances only. It does not perform MSA subsampling or MSA selection, and it does not predict orientations, as in the trRosetta method. As suggested in the trRosetta work [6] we anticipate that incorporating MSA subsampling and selection can further improve the precision of top L long-range contacts by around 16%.

We also investigated if our approach of splitting the input target sequence into 256-size crops was effective towards improving the precision of the structural domains in the case of longer protein targets (*L >* 256). Of all the CASP13 FM target domains we evaluated, we found that 14 of the domains had their corresponding target length greater than 256. For these 14 domains we asked what the change in precision would be (of the distance derived contacts) if we did not split the input sequence and built MSA for the entire sequence. We observed a drop in top L/5 long-range precision from 75.7% to 63.6% when the sequence splitting method was not used (see **Figure 6**). This is a clear boost of 20% in precision on average for the 14 domains. On the same 14 domains, it is worth noting that the average precision was 64.1% when alignments generated by the trRosetta method were used, i.e., a performance similar to the REALDIST version without sequence splitting. In other words, without sequence splitting, the MSAs from trRosetta method had a similar accuracy as our MSA generation method. These results suggest that if the technique of sequence splitting is integrated into existing methods for distance prediction (including trRosetta and DeepDist), a significant increase in accuracy may be observed. We further observed that this improvement in performance decreases as we evaluate a greater number of contacts. For example, when top L long-range contacts are evaluated, the improvement was only 9%.

**Figure 6:**
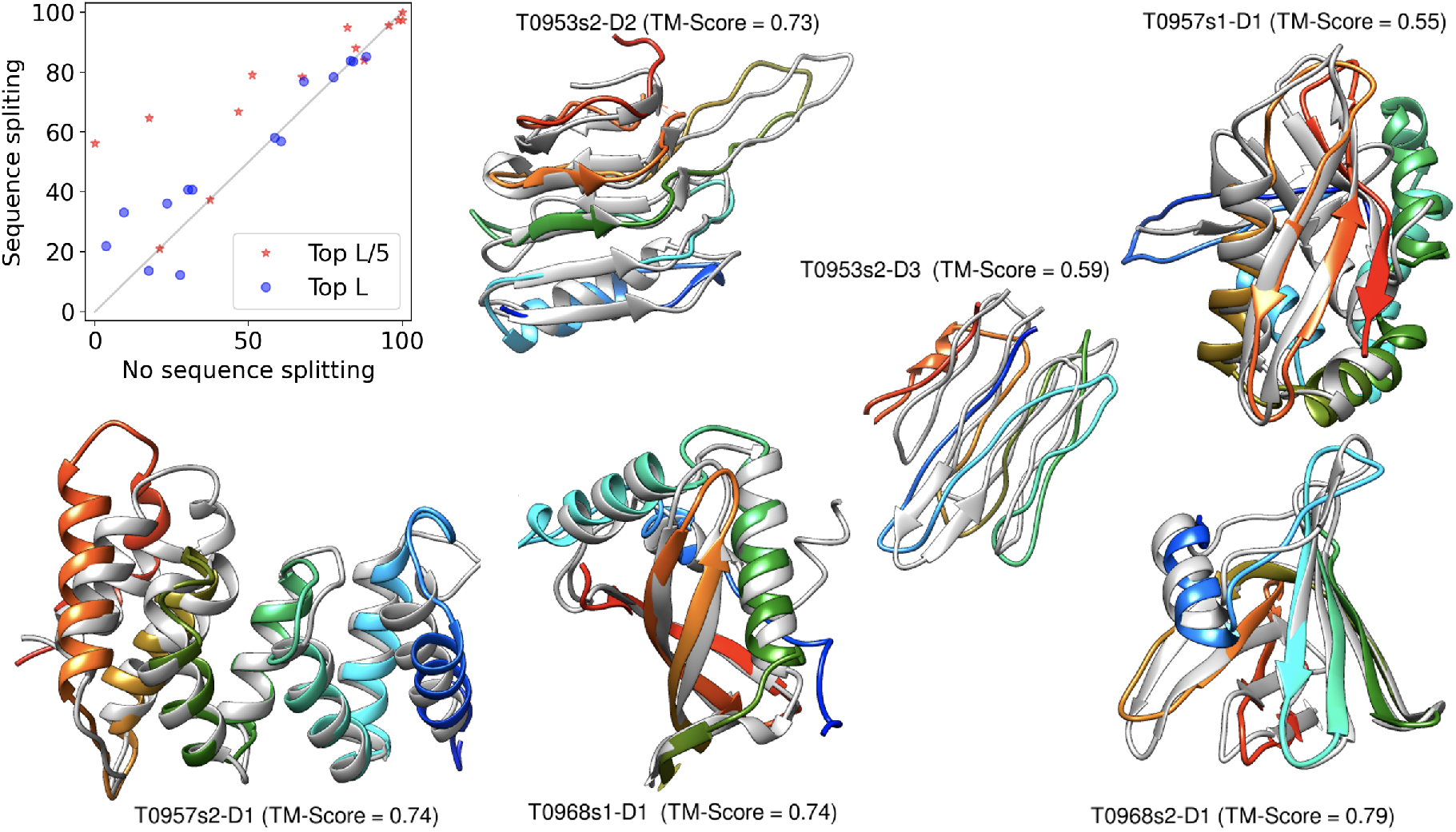
Improvement from multiple target sequence splitting observed by evaluating the precision of long-range distance-derived contacts on the 14 CASP13 FM domains with corresponding target length greater than 256 (scatter plot), and visualization of top one models built using Rosetta (with REALDIST distance constraints) superimposed with the native structure for CASP13 free-modeling domains whose true structures are publicly accessible.

### 3.4 Integrating the CATH and PISCES set for training

We observed that combining the PISCES set consisting of 28 thousand chains and the CATH set consisting of 25 thousand chains to form the development set results in around 2-5% increase in accuracy. We trained three ResNet models each trained using PISCES only, CATH only, and both CATH & PISCES as the development sets. Evaluation of these three models on the 131 CAMEO hard dataset and the 66 very hard subset reveals that using the PISCES dataset as a development set consistently but marginally outperforms using the CATH dataset (see **Table 3**). Combining the two datasets, on average, consistently yields slightly higher average precision than using any one of them. This gain, however, comes at almost double computational cost of training the ResNet.

**Table 3:** Gain from combining the PISCES and CATH datasets for deep learning evaluated on the 131 chains in the CAMEO set and for the subset of 66 very hard chains. Model predictions were performed without splitting the input sequences, i.e. the MSA-split technique was not used here.

| Dataset | Development set | $s \geq 24$ | | $s \geq 12$ | |
| --- | --- | --- | --- | --- | --- |
| | | $P_{L/5}$ | $P_L$ | $P_{L/5}$ | $P_L$ |
| 131 CAMEO hard | PISCES only | 79.0 | 55.9 | 80.8 | 65.15 |
|  | CATH only | 78.3 | 55.6 | 79.8 | 64.3 |
|  | CATH & PISCES | **81.1** | **57.6** | **83.3** | **66.8** |
| 66 CAMEO very hard | PISCES only | 70.0 | 45.9 | 71.0 | 55.2 |
|  | CATH only | 67.1 | 44.8 | 69.4 | 53.5 |
|  | CATH & PISCES | **72.1** | **47.7** | **74.8** | **57.1** |

### 3.5 Structure prediction using real-valued distances

Given that a true evaluation of predicted distances is to assess their utility towards building 3D models, we built models using the widely-used Rosetta *ab initio* protocol and evaluated the top models using the Template-Modeling score (TM-score) program [35]. Strictly following the CASP assessment practice of building 3D models for the entire targets, we ran the Rosetta *ab initio* program [36] for the CASP13 targets blindly, i.e., without any knowledge of the structural domains. Since running Rosetta requires a lot of CPUs, we built 3D models for all targets that were shorter than 250 residues. Also because of the limited number of CPUs available to us, we only generated 1000 models per target although it is recommended to generate 50 to 100 thousand decoys. We also only evaluated the top-one model selected using the overall Rosetta score (not the best of 1000). As shown in **Table 4**, the average TM-score of our method on these 17 domains is 0.57, which is similar to the performance of the top Human Group A7D [37] in the CASP13 competition. Our method, however, outperforms all the server group methods. As examples, for the targets whose structures are publicly available, our top models are superimposed with the native structure (see **Figure 6**). It is crucial to note that our method of converting predicted distances to Rosetta constraints is naive and empirical (see **Methods**) and improved techniques of translating distances to constraints and generating a much larger number of decoys will deliver significantly better models.

**Table 4:** Comparison of the evaluation of top-one models built using Rosetta (with REALDIST distance constraints) with the top Human Group (A7D) and the top Server Group (Quark) in the CASP13 challenge on free-modeling targets shorter than 250 residues.

| Method | TM-Score | RMSD | GDT-TS |
| --- | --- | --- | --- |
| Group #43 (A7D) | 0.56 | 9.35 | 50.84 |
| Group #145 (Quark) | 0.49 | 8.36 | 43.45 |
| REALDIST + Rosetta | **0.57** | **7.11** | **51.23** |

To further our comparison with trRosetta, we chose to build 3D models for the chain H of the protein target ‘5WB4’ in the CAMEO dataset—the target for which the trRosetta method was demonstrated to deliver a remarkable accuracy. For this target we ran REALDIST to predict real-valued distances and translated the distances into Rosetta modeling restraints. As shown in **Figure 3B**, REALDIST distance predictions for this protein are remarkably accurate. Next, 1000 models were built using the Rosetta ab initio protocol and the model with the lowest energy score was selected as the top predicted model (see **Methods**). Evaluation of this top model using the TM-score program demonstrated an extremely accurate model with a TM-score of 0.94 (see **Figure 3C**). Notably, TM-scores of the template-based models by HHpred, IntFOLD5-TS, and Raptor-X were around 0.4 (in the CAMEO competition), and the TM-scores by Robetta and trRosetta were 0.879 and 0.921 respectively [6]. It is also worth noting that REALDIST and trRosetta predictions were not truly blind, i.e., predictions were made after the true structure was released.

## 4 DISCUSSION

We acknowledge that our method has limitations; however, these limitations can be addressed. First, unlike the trRosetta method, REALDIST does not predict dihedral angles (orientations) or secondary structures. Since the focus of this work is to demonstrate the sole potential of real-valued distances, we intentionally skipped these auxiliary predictions. Although this is currently a limitation, REALDIST can be upgraded to predict orientations and secondary structures. Second, our method of alignment generation does not generate a single MSA for a protein that is longer than around 256 residues. Third, during training, we do not use any kind of MSA augmentation, such as changing MSA information so that the model sees different information each time. A limited amount of available solid-state disks (SSDs) and graphical processing units (GPUs) kept us from performing more thorough augmentations. Finally, our real-valued distance prediction models do not predict any kind of confidence or probabilities associated with the predicted distance values. One technique to address this limitation is to predict error bounds (standard deviations) for each distance prediction. An appropriate loss function must be designed for this purpose.

## 5 CONCLUSION

Our real-valued distance prediction method demonstrates state-of-the-art results and unveils a promising new direction in the field of protein structure prediction. The utility of the predicted distances was also demonstrated through distance-guided three-dimensional structure prediction. It is exciting that a standard ResNet model, that only predicts a real-valued distance map, can perform on par or better than the state-of-the-art methods for distance prediction. This real-valued distance prediction approach offers a new direction.

## 6 AVAILABILITY

REALDIST’s source code and trained models are publicly available at https://github.com/ba-lab/realdist/.

## 7 ACKNOWLEDGEMENTS

We acknowledge financial support from the US National Science Foundation to B.A. (award number 1948117). We are also thankful to Dr. Chengxin Zhang at University of Michigan and Dr. Jie Hou at Saint Louis University for providing numerous suggestions to improve the manuscript.

---

**Table S1.** Metagenomic sequence databases used for running DeepMSA.

| Database | Size* | Download location |
| --- | --- | --- |
| Soil Reference Catalog | 450GB | http://wwwuser.gwdg.de/~compbiol/plass/ |
| Marine Eukaryotic Ref. Catalog | 70GB | http://wwwuser.gwdg.de/~compbiol/plass/ |
| MetaClust | 350GB | https://metaclust.mmseqs.com/ |
| Peptide DB | 150GB | http://ftp.ebi.ac.uk/pub/databases/metagenomics/peptide_database/ |

\*Size is approximate and includes the uncompressed fasta file and the index (.ssi) file.

